# DeepSVP: Integration of genotype and phenotype for structural variant prioritization using deep learning

**DOI:** 10.1101/2021.01.28.428557

**Authors:** Azza Althagafi, Lamia Alsubaie, Nagarajan Kathiresan, Katsuhiko Mineta, Taghrid Aloraini, Fuad Almutairi, Majid Alfadhel, Takashi Gojobori, Ahmad Alfares, Robert Hoehndorf

## Abstract

**Motivation:** Structural genomic variants account for much of human variability and are involved in several diseases. Structural variants are complex and may affect coding regions of multiple genes, or affect the functions of genomic regions in different ways from single nucleotide variants. Interpreting the phenotypic consequences of structural variants relies on information about gene functions, haploinsufficiency or triplosensitivity, and other genomic features. Phenotype-based methods to identifying variants that are involved in genetic diseases combine molecular features with prior knowledge about the phenotypic consequences of altering gene functions. While phenotype-based methods have been applied successfully to single nucleotide variants as well as short insertions and deletions, the complexity of structural variants makes it more challenging to link them to phenotypes. Furthermore, structural variants can affect a large number of coding regions, and phenotype information may not be available for all of them.

**Results:** We developed DeepSVP, a computational method to prioritize structural variants involved in genetic diseases by combining genomic and gene functions information. We incorporate phenotypes linked to genes, functions of gene products, gene expression in individual celltypes, and anatomical sites of expression, and systematically relate them to their phenotypic consequences through ontologies and machine learning. DeepSVP significantly improves the success rate of finding causative variants in several benchmarks and can identify novel pathogenic structural variants in consanguineous families.

**Availability:** https://github.com/bio-ontology-research-group/DeepSVP

## 1 Introduction

Structural genomic variants are genomic variants that affect more than 50 base pairs and include copy number variants, insertions, and deletions (Eichler, 2019). Many structural variants (SVs) are implicated in heritable diseases (Sudmant *et al.*, 2015). While there have been several efforts to predict and prioritize pathogenic genomic variants (Eilbeck *et al.*, 2017), predicting the functional impact of SVs discovered through genome sequencing studies remains challenging due to the diversity of variant size and type; SVs may cover multiple coding and non-coding regions, overlap several genes, and are affected by haploinsufficiency and triplosensitivity (Kidd *et al.*, 2008). Methods for predicting the pathogenicity of genomic variants may be based on their impact on protein structure, measures of sequence conservation, or function (Eilbeck *et al.*, 2017). However, due to the complexity of the SVs, including the variant size, type, and overlap with multiple genes, designing methods that determine structural variant pathogenicity is more challenging. Several efforts for analyzing the clinical impact of structural variants have focused on well-matched cases and controls. For instance, by evaluating the loci and the respective pathways that may be impacted by a structural variant at these loci, it became possible to define novel genes involved in complex disorders such as autism (Pinto *et al.*, 2010) or immune-related disorders (Rossin *et al.*, 2011). While there are several methods to identify disease-associated variants in cohorts, it is more challenging to discover disease-associated variants that exist in a single sample or pedigree, in particular in rare Mendelian disorders (Sanchis-Juan *et al.*, 2018).

Methods that evaluate the functional consequence of structural variants in individual genomes use different strategies. Several approaches include genomic information, such as variant length, haploinsufficiency measures, or GC contents, to separate pathogenic from benign SVs (Hehir-Kwa *et al.*, 2010; Sharo *et al.*, 2020). Furthermore, the predicted pathogenicity of deleterious single nucleotide variants within a structural variant can be used to estimate pathogenicity of SVs (Ganel *et al.*, 2017). Additionally, phenotypes associated with a loss of function in single genes has also been used for prioritizing SVs (Doelken *et al.*, 2012; Köhler *et al.*, 2014).

Phenotype-driven variant prioritization methods aim to link variants to the phenotypes observed in individuals using prior knowledge (Eilbeck *et al.*, 2017). Commonly, the link is established using a similarity measure between phenotypes associated with a variant or gene and the phenotypes observed in a patient (Smedley *et al.*, 2015). Phenotype-based methods are successful in finding disease-associated variants (Shefchek *et al.*, 2020) but suffer from the limited information about variant– or gene–phenotype associations. One way to overcome this limitation is to utilize phenotypes observed in model organisms and link them to human phenotypes (Smedley *et al.*, 2013; Shefchek *et al.*, 2020); however, even when including phenotypes from model organisms a large portion of human protein-coding genes remain without associations, thereby limiting the success of phenotype-based methods to variants or genes that have previously been studied either in humans or animal models, or relying on guilt-by-associations approaches in which information about phenotypes is propagated through associations such as interaction networks (Smedley *et al.*, 2014).

Several deep learning methods are now available that can predict phenotypes (Zhou *et al.*, 2019; Kulmanov and Hoehndorf, 2020) or associate phenotypes with different types of information available for genes, including functions of gene products and site of expression (Chen *et al.*, 2020; Smaili *et al.*, 2019). These methods use machine learning to relate information through background knowledge contained in ontologies, and can accurately identify phenotype-associated genes without prior knowledge about phenotypes, often significantly improving over the use of semantic similarity measures (Kulmanov *et al.*, 2020). A limitation of these methods is that they are often transductive instead of inductive (Kulmanov *et al.*, 2020), i.e., the diseases or disorders for which associated genes are predicted should already be available at the time of training the model. As these methods require information about disease-associated phenotypes during training, they will therefore not generalize to entirely new cases, thereby limiting their application in identifying phenotype-associated genomic variants.

We developed a machine learning method that predicts whether a copy number variant (i.e., a structural variant that is either a duplication or deletion) is pathogenic and involved in the development of specific phenotypes. Our method combines genomic information and clinical phenotypes, and leverages a large amount of background knowledge from human and animal models; for this purpose, we extend an ontology-based deep learning method to allow inductive inference. We demonstrate that our method improves over the state of the art in detecting pathogenic deletions or duplications, in particular improving the precision in finding phenotype-associated variants. We further apply our method to the diagnosis of a family with congenital disease involving infantile spasms and seizures for which previous analysis of single nucleotide variants in whole exome and whole genome sequencing data found no associated variant. We make DeepSVP freely available as a Python package at https://github.com/bio-ontology-research-group/DeepSVP.

## 2 Materials and Methods

### 2.1 Inputs, outputs, and problem statement

DeepSVP is a machine learning model that takes as input a set of SVs in Variant Call Format (VCF) format together with a set of phenotypes encoded using the Human Phenotype Ontology (HPO). DeepSVP outputs a list of the variants from the input VCF file ranked by their probability of being associated with (or causative of) the set of phenotypes provided as input.

### 2.2 Data sources and ontologies

We use as training and testing dataset the set of pathogenic and benign SVs aligned to the human reference genome GRCh38 obtained from the database of genomic structural variation (dbVar) (Griffith and Griffith, 2004) downloaded on 8th Feb 2020. We use functional and phenotypic characteristics for genes in the human genome, in particular the phenotypes associated with human genes in the HPO database (Köhler *et al.*, 2019), the phenotypes associated with mouse orthologs in the Mouse Genome Informatics (MGI) database (Bult *et al.*, 2019), the functions of gene products from UniProt (UniProt Consortium, 2019), gene expression in human celltypes (Tabula Muris Consortium *et al.*, 2018) and the anatomical site of expression from the GTEx tissue expression database (GTEx Consortium, 2015). These annotations are characterized using the HPO (Köhler *et al.*, 2019), Mammalian Phenotype Ontology (MP) (Bult *et al.*, 2019), Gene Ontology (GO) (Gene Ontology Consortium, 2019), Cell Ontology (CL) (Diehl *et al.*, 2016), and the anatomical site of expression (UBERON) ontology (Mungall *et al.*, 2012). Detailed information about the data sources and ontologies is provided in the Supplementary Materials Section 1.

We annotate SVs with a set of genomic features using public databases. We use AnnotSV v2.3 (Geoffroy *et al.*, 2018), which uses data from multiple external databases to annotate and rank SVs. A detailed description of variant-based features is provided in the Supplementary Materials Section 2.

### 2.3 Embedding ontology-based features

We use the DL2Vec (Chen *et al.*, 2020) method to encode ontology-based features associated with genes in a low-dimensional feature vector. DL2Vec “embeds” ontologies and their annotations in a real-valued vector space. For this purpose, DL2Vec first generates a graph *G* = (*V, E*) from the ontology axioms in which nodes *V* represent classes or entities annotated with ontology classes, and edges *E* represent axioms that hold between these classes (Chen *et al.*, 2020). DL2Vec then explores the graph using random walks, and generates embeddings from these walks using Word2Vec. We apply DL2Vec to the annotations and ontologies for the genes associated with HPO, GO, MP, CL, and UBERON. As each feature is available for a different number of genes, we generate embedding for each kind of feature separately. As parameters for DL2Vec, we use 100 random walks with a walk length of 25; we use the Word2Vec (Mikolov *et al.*, 2013) skip-gram model to generate the embeddings from the walks with 10 window size, 1 as the minimum count value, and an embedding size of 100. We train the skip-gram model for 20 epochs. As a result, we obtain a real-valued feature vector for each gene (and ontology class) of size 100.

We encoded the input set of phenotypes using DL2Vec to generate an embedding of the patient phenotypes. As there will not likely be a representation of an entity associated with all phenotypes used as input to our method, we first update the graph used by DL2Vec to add a new node for the patient; this new node is associated with all phenotypes used as input. We then generate a new embedding for this node by performing random walks and then updating the pre-trained skip-gram model using these walks. This approach allows the skip-gram model to generate an embedding for a new patient (specified entirely by a set of phenotypes) while considering the full DL2Vec graph generated from the ontology.

### 2.4 Estimating variant pathogenicity by supervised prediction

We build a machine learning model that ranks SVs depending on their predicted pathogenicity and the relations between genes affected by the SV and the phenotype observed in affected individuals. Additionally, our approach consider several genomic features of each SVs, such as the coding sequence length overlapping with the SV, GC content, and haploinsufficiency and triplosensitivity scores to measure the dosage-sensitivity for genes/regions. Using the VCF file of the patient and HPO-encoded set of phenotypes, all features are generated by annotating the VCF file. Figure 1 presents a high-level summary of training our model. The DeepSVP prediction model consists of two parts: a phenotype prediction model based on matching the gene features with the phenotypes observed in the affected individuals (Figure 1-b); and a combined prediction model based on the phenotype prediction scores and the genomic features of the variant (Figure 1-c). A detailed description of the model and training process is provided as Supplementary Materials Section 3.

**Fig. 1:**
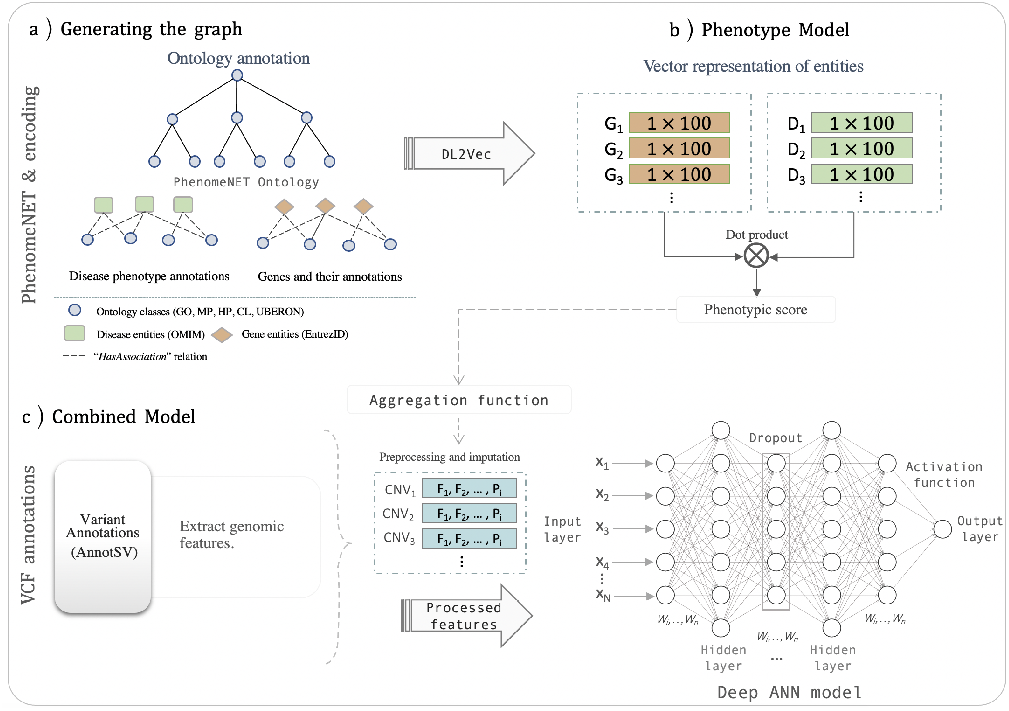
Overview over DeepSVP training model. (**a**) Generate a graph from the ontology axioms in which nodes represent classes or entities annotated with ontology classes, and edges represent axioms that hold between these classes. (**b**) The DL2Vec workflow takes a set of phenotypes as input and predicts whether a gene is likely associated with these phenotypes using several types of background knowledge. (**c**) The combined model uses the prediction score of the DL2Vec model combined with genomic features derived from variants; we performed two aggregation functions for the genes within the variants (maximum and average). We trained the model and output a prediction score for each variant that determining how likely the variant is causative of the disease in our training set of structural variants. Abbreviations: `G:` Genes, `D:` Diseases, `F:` Features, `P:` Phenotypic score.

### 2.5 Predicting disease-associated structural variants

Our prioritization algorithm relies on finding the “causal” variants which are (a) pathogenic and (b) involved in developing a set of phenotypes observed in a patient. We investigated the performance of our machine learning classifier to predict the causative structural variant based on genomic characteristics of the variant, the functions and phenotypes that are likely altered by the variant, and the phenotypes observed in the individual in which the variant was detected. In our experiments, we considered as positive instances all the causative variants in our training set with the disease phenotypes for which they are causative. We separate the positive instances from two types of negatives instances, the benign variants which do not change any protein function and are not implicated in a disease, and pathogenic (but non-causative) variants that may be pathogenic but are not related to the disease phenotypes observed in a patient (but potentially related to another set of phenotypes).

We randomly split our dataset by diseases and associated variants into 85% for training and 15% for testing, where the set of diseases in training are different from the testing (to reflect the application of our method to entirely new diseases that have not been used for training). We use 15% of the training set as a validation set. We used the training and validation sets to train and tune model hyperparameters and select the best models, while the test set for reporting the evaluation metrics.

The number of negative samples (representing the set of variant-phenotype pairs that are either benign or the pathogenic but not causative variants) is significantly higher than the number of positives samples (representing the causative variants of a particular disease). For that reason, we randomly sample 5 times from the negative set a number of instances equal to the positive instances and use these as negatives for training.

We build a separate model for each ontology dataset and aggregation operation (either maximum or average). The architecture of the prediction model was derived by hyperparameter optimization using Scikit-Optimize with Bayesian optimization (`skopt`) repeated 15 times. We provide a detailed description of the set of hyperparameters we tuned and use in the Supplementary Materials Section 4. We implemented our models using Keras with a TensorFlow backend, and training was performed on a single Nvidia Tesla V100 GPU. We evaluated the performances of our model in the testing set using the Area Under the Receiver Operating Characteristics Curve (ROCAUC), F1-score, the Area Under the Precision-Recall Curve (PRAUC), and the Diagnostic Odds Ratio (DOR) (Glas *et al.*, 2003).

### 2.6 Benchmarking of SV prioritization methods

We compared the performance of DeepSVP to two related methods that can rank or classify SVs, StrVCTVRE (Sharo *et al.*, 2020), and AnnotSV version 2.3 (Geoffroy *et al.*, 2018). StrVCTVRE is a structural variant impact predictor that captures a set of genomic features for structural variants relating to the conservation, gene importance, coding region, expression, and exon structure, trained using a random forest classifier. AnnotSV provides a classification for each SV based on recommendations for the interpretation of copy number variants (Riggs *et al.*, 2019) and classifies variants into pathogenic, likely pathogenic, of uncertain significance, likely benign, and benign. AnnotSV can use the phenotype-based method Exomiser (Smedley *et al.*, 2015) to determine whether phenotypes are consistent with previously reported cases, and incorporate the phenotype-based score in the variant classification process. We rank variants based on the class assigned by AnnotSV, descending from pathogenic to benign.

As a benchmark set, we use the disease-associated variants added to the dbVar database between 8 February 2020 and 2 July 2020. Our training dataset and that of StrVCTVRE and AnnotSV are limited to the set of variants that have been added to dbVar prior to that date. 1,503 disease-associated variants were added between 8 February 2020 and 2 July 2020, covering 579 distinct diseases, 175 of which are not linked with any disease in our training set. The evaluation using entirely unseen disease-associated variants helps us to estimate how well DeepSVP can prioritize novel variants under more realistic conditions. We created synthetic patient samples by inserting a single causative variant into a whole genome sequence from the 1000 Genomes Project for structural variant (1000 Genomes Project Consortium *et al.*, 2012). The set of SVs in 1000 Genomes contains a total of 68,697 variants for 2,504 individuals from 26 populations. Using the 1000 Genomes frequencies for all populations, we exclude all variants with Minor Allele Frequency (MAF) of more than 1% which results in 2,391 variants remaining. Then we link the phenotypes associated with (OMIM) that are associated with the causative variant in dbVar (using `phenotype_annotation.tab`) to the synthetic genome. We consider the combination of the synthetic genome and HPO phenotypes as a synthetic patient sample. We repeat this for all 1,503 causative variants.

### 2.7 Whole genome sequencing and SV calling

We collected blood samples for a Saudi family consisting of five individuals, two unaffected parents, two affected children and one unaffected child. We performed whole genome shotgun sequencing on all individuals. Details for alignment and structural variant calling are provided in the Supplementary Materials Section 5.

### 2.8 Ethical approvals

This study was approved by the Institutional Research Board of the King Abdullah International Medical Research Center (RC 16/113 and RC16/211/R2), and the Institutional Bioethics Committee at King Abdullah University of Science and Technology (17IBEC08_Gojobori). All patients have been consented to be enrolled in this study, a written consent form was obtained from all subjects or their parents or legal guardians in the case of minors who are aged 16 years old or younger.

## 3 Results

### 3.1 DeepSVP predicts phenotype-associated variants in synthetic genomes

We developed DeepSVP as a method to identify phenotype-associated SVs (deletions and duplications) for patients based on personal genomic data and the phenotypes observed in a patient. The aim of the DeepSVP model is not only to detect potentially pathogenic SVs, but identify the variants that are “causative” of a set of phenotypes observed in a patient. We consider a variant as cause of a set of phenotypes when it is both pathogenic (i.e., disrupts the normal functioning of one or more genes) and contributes to the development of the phenotypes. This approach is motivated by the observation that even healthy individuals may have pathogenic or potentially pathogenic variants that do not result in abnormal phenotypes. Therefore, detecting pathogenicity of a variant alone is typically not sufficient to establish causality (MacArthur *et al.*, 2014). Our workflow uses as input a set of variants and a set of phenotypes observed in an individual; for each variant–phenotypes pair, DeepSVP predicts whether the variant is likely to be causative of the phenotypes, and outputs a ranked list of variants. DeepSVP uses the phenotypes arising from a loss of function in mouse, phenotypes associated with human genes, the anatomical site of gene expression, gene functions, and celltypes in which genes are expressed, as background knowledge, and links these to the abnormal phenotypes observed in the individual in which the structural was detected. To make the predictions based on these different features types, we embed them into a shared representation space using a feature learning method applied to ontologies (Chen *et al.*, 2020). We then combine the resulting embeddings with sequence-derived features that can be used to predict the pathogenicity of a variant, and use a neural network model to predict whether a variant is associated with patient phenotypes. The workflow for training the DeepSVP model is illustrated in Figure 1, and the workflow for inference in Supplementary Figure 7.

We used 85% of the diseases in our dataset, together with all their associated structural variants, for training and 15% for testing. We first evaluate DeepSVP’s performance in finding disease-causing structural variants on these 15%. Supplementary Table 2 and Supplementary Figure 3 show a summary of DeepSVP’s performance. We find that the our model can separate positive from negative cases with a ROCAUC ranging from 0.8913 (using only celltype of expression as background knowledge) to 0.9534 (combining all background knowledge). However, the evaluation on a testing dataset that resembles the training data will not be indicative of the performance of the model in a realistic setting where the aim is to identify a single disease-associated variant among potentially hundreds or thousands of candidates within a genome.

As a more realistic evaluation of our model, we generate synthetic patient data in which we combine the variants from the genome sequences in the 1000 Genomes project, insert a single disease-causing pathogenic variant, and associate this synthetic genome with the phenotypes of the variant. We then apply our model to all SVs in this synthetic patient, rank the resulting variants based on DeepSVP’s prediction score, and evaluate the results. For this evaluation, we select an independent dataset of disease-associated SVs, i.e., the set of variant–disease pairs added to dbVar after we obtained training data for our model. Our evaluation set contains 1,503 variants associated with 579 distinct diseases in OMIM and overlapping with 1,926 unique genes. There are 175 diseases (associated with 640 variants) that were not present in our training data. We create synthetic patient samples for all variant–phenotype pairs in this evaluation set. We also compare the results of our model with another method for identifying disease-associated variants, StrVCTVRE, and AnnotSV. Table 1 and Supplementary Figure 4, shows the amount of disease-associated variants we identify at different ranks. We evaluate the recall separately for variants that are associated with a disease already present in our training data and variants that were not.

**Table 1.**
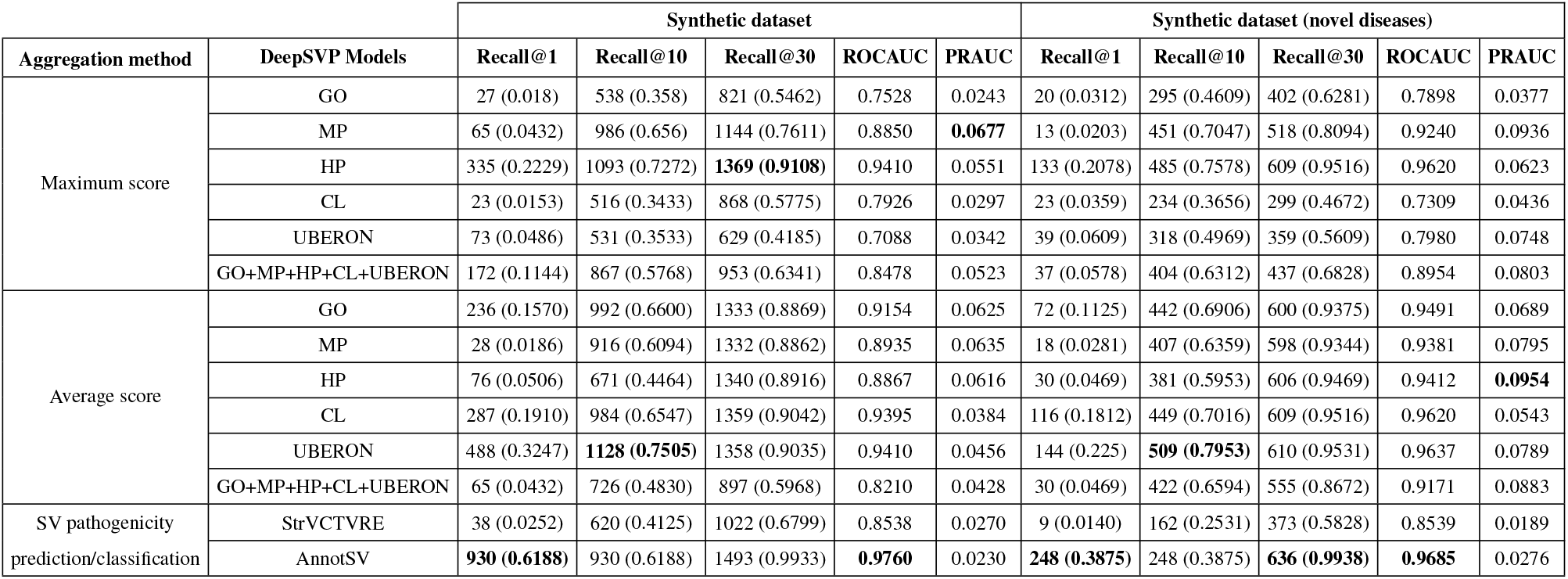
Summary of the evaluation for predicting causative variants in the benchmark dataset of dbVar, time-based split, for 1,503 newly added variants along with the evaluation for 640 newly added variants associated with 175 new diseases which were not present in our training dataset. The evaluation inserts one disease-associated structural variant in a whole-genome and reports the rank at which the inserted variant was recovered. We report the absolute number of variants recovered at each rank. AnnotSV classifies variants in five categories, and ranks for AnnotSV are computed based on ranking variants from “pathogenic” to “benign”; variants with the same classification are tied and we report all variants at the same (best) rank; only AnnotSV predictions produce ties between variants. Best performing results for each measure are indicated in bold.

The performance using different DeepSVP models varies depending on the availability of the features as well as the gene annotations that are available for each type of feature. The benchmark dataset (i.e., the list of variants we inserted) covers 1,920 unique genes, with the largest variant containing 129 genes and the smallest variant containing one gene. There are 635, 963, 1214, 1360, 493 missing annotations from these genes for features represented using GO, MP, HP, CL, and UBERON, respectively. However, the models are still able to predict causative variants using the annotations of the remaining genes (1285, 957, 706, 560, 1427 for GO, MP, HP, CL, and UBERON) which have an annotation and a corresponding “representation”. The evaluation demonstrates that DeepSVP can identify the variants in novel (i.e., unseen during training) disease-associated variants with one, or more than one, gene and still improve recall at rank one over some established methods. We also find that DeepSVP significantly improves prioritization of disease-associated variants over StrVCTVRE (*p* < 3.9 × 10^−13^, Mann-Whitney U test), a method that uses similar features as DeepSVP to determine pathogenicity of variants but does not rely on information about phenotypic or functional consequences. We further evaluated the classifications provided by AnnotSV; AnnotSV classifies variants into five classes (pathogenic, likely pathogenic, unknown significance, likely benign, benign) which we rank by sorting variants based on this classification (from pathoenic variants as highest to benign variants as lowest); using this ranking, AnnotSV has the highest recall at rank 1 (i.e., variants predicted as “pathogenic”) but it also predicts multiple variants at the same rank (i.e., as pathogenic variants), leading to a lower precision compared to DeepSVP or StrVCTVRE which rank variants without ties.

While we are able to identify disease-associated SVs using phenotype information, the phenotypes reported with a patient-derived sample will not always be complete or as comprehensive as in our evaluation. To determine the effect of different phenotype associations, we further evaluated the performance of DeepSVP when only partial phenotype data is available. We repeat our experiment using synthetic patient samples while randomly removing between 10% and 50% of the phenotypes associated with the sample (Supplementary Tables 3, 4, 5, and 6). We find that even when reducing the number of associated phenotypes to 50%, the performance of our model remains comparable; however, when removing phenotype information entirely using the combined model, the predictive performance drops compared to a model that includes information about phenotypes, which results in 27 (1.8%) of causative variants ranked first and 575 (38.3%) in the top 10.

### 3.2 DeepSVP identifies a disease-associated variant in a consanguineous family

We applied DeepSVP to investigate whole genome sequencing data in nine consanguineous families from Saudi Arabia (Alfares *et al.*, 2020) where the clinical presentation is suggestive of genetics underlying etiology with two or more affected individuals, and all previous genetic analyses based on clinical and whole exome analysis were negative.

First, we investigated single nucleotide variants and small insertions and deletions as possible explanation of the underlying genetics etiology. However, we were unable to identify a possible disease-associated variant in all nine families using this approach (Alfares *et al.*, 2020).

We applied DeepSVP to rank structural variants for possible explanation of the phenotype. In one family, the affected individuals showed hypotonia (`HP:0001290`), developmental delay (`HP:0001263`), infantile spasms (`HP:0012469`), strabismus (`HP:0000486`), and seizures (`HP:0001250`); after removing all common variations, we ended up with 47 SVs requiring further investigation. DeepSVP ranked one duplication in chromosome `2q24.3(NC_000002.12: g.164062341_166264282dup)` as top rank using the combined model based on the maximum score (see Supplementary Table 7 for other DeepSVP models); this duplication contains 16 genes (*COBLL1*, *CSRNP3*, *GALNT3*, *GRB14*, *LOC100506124*, *LOC101929633*, *LOC102724058*, *SCN1A*, *SCN1A-AS1*, *SCN2A*, *SCN3A*, *SCN9A*, *SLC38A11*, *SNORA70F*, *TTC21B*, *TTC21B-AS1*). Duplication of 2q24.3, including the cluster of voltage-gated sodium channel genes, is linked with hypotonia, seizures, and neonatal epilepsy in several unrelated cases (Simonetti *et al.*, 2012; Okumura *et al.*, 2011; Firth *et al.*, 2009). We further confirmed the variant with length 2201941 using Array Comparative Genomic Hybridization in a clinical laboratory. DeepSVP outputs the gene based on which phenotypic similarity was established (using the maximum phenotype score), and among the 16 genes, the *SCN1A* gene has the maximum phenotype score using MP model. The heterozygote loss of function of *SCN1A* in mouse is a model of Dravet syndrome (Kim *et al.*, 2018; Miller *et al.*, 2013) and resembles many of the phenotypes observed in the family we analyzed. Our results shows that DeepSVP can not only identify disease-associated SVs but further yield interpretable results that can provide actionable clinical information. We compared the results with predictions of AnnotSV and StrVCRTRE. AnnotSV classified 11 variants out of 47 as pathogenic, including the variant we identified. StrVCRTRE scored only 6 variants out of 47 (the remaining variants are either less than 50*bp* in length or not exonic, and thereby out of the scope of StrVCRTRE), and the variants we identified ranked 4 out of 6.

## 4 Discussion

### 4.1 Related work

A seminal study investigated the application of phenotype-similarity in CNVs (Doelken *et al.*, 2012) and the results let to the PhenoGramViz method and tool (Köhler *et al.*, 2014). PhenoGramViz, as well as AnnotSV, use phenotypes to rank and prioritize SVs using phenotype information. To overcome the limitation of missing information about phenotypes, PhenoGramViz and AnnotSV rely on mouse phenotypes. While mouse phenotypes increase the coverage of genes with phenotype associations, there are nevertheless a large number of genes for which no phenotype associations are available. DeepSVP overcomes the limitation of missing phenotypes by incorporating information related to genes through ontologies, mainly the functions of gene products, gene expression in individual celltypes, and anatomical sites of expression and systematically relating them to their phenotypic consequences through ontologies. The phenotype-based prediction model in DeepSVP is modular and can be utilized as part of other methods such as AnnotSV.

PhenoGramViz is not a method that directly prioritizes structural variants but relies on visualizing ranking results and exploration by users. While this is useful in targeted studies, DeepSVP can be applied as a component of computational workflows while still enabling interpretation of results. AnnotSV provides a classification rank for each SV using five classes based on their overlap with known variants from different data sources, and aims to implement clinical classification guidelines for variants. DeepSVP, on the other hand, provides pathogenicity prediction for each variant rather than categorize them and includes phenotype prediction models not only to identify relatedness to known phenotypes but also to predict new ones; it may therefore be more suitable for generating hypotheses about phenotype-associations of SVs that do not overlap with known disease genes. StrVCRTRE is a method that also directly predicts pathogenicity of SVs and uses similar features related to the gene importance, coding sequence, and expression, which allows us to compare directly. A key difference between DeepSVP and StrVCRTRE is DeepSVP’s use of phenotype information while StrVCRTRE does not rely on phenotype information which improves prediction results significantly. Furthermore, StrVCRTRE ranks only the exonic variants, while DeepSVP ranks both exonic and intronic based on the availability of the genomics features.

We evaluated and compared DeepSVP with AnnotSV and find that DeepSVP does not improve over AnnotSV with respect to recall but it improves the precision of finding phenotype-associated variants; AnnotSV is not a method to rank variants but rather to classify variants in categories, which may lead to multiple variants being classified as “pathogenic” and therefore decrease precision. DeepSVP ranks variants without ties and shows generally higher precision in our evaluation; it may also allow DeepSVP to more easily be applied in computational workflows where high precision is desirable. Since version 3.0, AnnotSV also provides a more fine-grained ranking of variants from which the variant classification is derived; we did not compare DeepSVP with this version of AnnotSV as it was released after our synthetic, time-based dataset was created. In the future, we intend to provide additional comparisons of DeepSVP with the different outputs of AnnotSV.

### 4.2 Machine learning with semantic background knowledge: from transductive to inductive

DeepSVP relies on machine learning for predicting pathogenic and phenotype-associated with the patient. For this purpose, it relies on advances in machine learning with ontologies that incorporate the background knowledge contained in ontologies in the form of axioms and annotations to ontology classes (Kulmanov *et al.*, 2020). Many such approaches convert ontologies into a graph-based form based on syntactic patterns within the ontology axioms and then apply a graph embedding on the resulting graph (Kulmanov *et al.*, 2020). In DeepSVP, we use DL2Vec which includes a large variety of ontology axioms and can significantly improve the phenotype–based prediction of disease genes. While these ontology-based methods rank genes, DeepSVP directly ranks SVs based on the genomic and phenotypic features collected from public databases, and the phenotypes observed in a patient. We precomputed the embeddings for genes based on different features (function, phenotype, and expression in celltypes and anatomical parts).

Furthermore, we extended the ontology-based machine learning methods to an inductive setting where we can predict associations between genes and patients that are defined by their phenotypes which are not known at the time of training DeepSVP. We also applied a rank-based normalization, similar to the method applied by PhenoRank (Cornish *et al.*, 2018), and use the resulting score instead of prediction scores of the neural network model; this transformation is useful when predicting relations where one argument remains fixed as it projects prediction scores into the same distribution.

While we implemented a two-step approach in which we first predict associations between genes and patient phenotpyes, and second the pathogenicity and phenotypic relatedness of the variant to the patient phenotypes, it may also be possible to design a model that is trained in an end-to-end fashion in the future. The challenge is the potentially open-ended number of genes to consider.

### 4.3 Clinical application and utility

We evaluated the performance of DeepSVP on a series of real genomes from Saudi individuals where the clinical presentation is suggestive of genetic diseases to assess how well we could recover potentially pathological variants in genes already associated with the disease. We used the whole genome sequencing data from all family members, and we apply family filtering according to the suitable inheritance pattern. We applied DeepSVP to rank SVs for a possible explanation of the phenotype. In one family, our model was able to find the causative variants associated with the patient phenotypes using the combined prediction model that integrates all the phenotypes information, and also to highlight a candidate gene underlying the main phenotypic manifestations. AnnotSV also classified this variant as pathogenic, together with 10 other variants.

We implemented two models for aggregating phenotypic relatedness between genes and patient phenotypes, one using the maximum and another using the average of scores of all genes. These correspond to two different mechanisms through which a structural variant elicits abnormal phenotypes: the maximum model is applicable when a single gene within the variant is (primarily) causative for the phenotypes, whereas the average model is applicable in the oligogenic case when multiple genes affected by a structural variant are causative and may contribute different pathologies.

We make DeepSVP freely available to use as a free software commandline tool, including all the steps to train the model. DeepSVP uses as input an annotated VCF file of an individual and clinical phenotypes encoded using HPO. DeepSVP can be used as a part of interpretation workflows in a clinical setting, or incorporated in interactive variant exploration methods.

## Acknowledgements

This work was supported by funding from King Abdullah University of Science and Technology (KAUST) Office of Sponsored Research (OSR) under Award No. URF/1/3790-01-01, FCC/1/1976-08-01,and FCC/1/1976-08-08. We acknowledge support from the KAUST Supercomputing Laboratory.

## Data availability

All data underlying this article except sequencing data are freely available at https://github.com/bio-ontology-research-group/DeepSVP. Sequencing data for the Saudi family cannot be shared publicly to protect the privacy of participating individuals. Requests for data access can be submitted to the Institutional Research Board of the King Abdullah International Medical Research Center and the Institutional Bioethics Committee at King Abdullah University of Science and Technology.

