## Supplemental materials for "DeepSVP: Integration of genotype and phenotype for structural variant prioritization using deep learning"

#### 1 Data sources and ontologies

The training dataset of variants contains 14,197 (10,401 deletions, 3,796 duplications) pathogenic or likely pathogenic structural variants and 4,477 variants associated with one or more diseases (3,737 deletions, 586 duplications), as well as 25,890 (13,742 deletions, 12,148 duplications) benign or likely benign structural variants.

For each pathogenic structural variant, we defined variant–disease pairs with associated diseases from Online Mendelian Inheritance in Men (OMIM) database [1]. There are 3,805 structural variants linked with one or more than one OMIM disease; if a variant is associated with *n* OMIM diseases, we generate *n* variant–disease pairs. As a result, we obtained 5,907 causative pathogenic variant–disease pairs. As negative training pairs we selected both benign and pathogenic variants and associate them with a randomly selected disease; we include pathogenic variants in the negative training pairs to simulate the case where variants may be pathogenic but not associated with the phenotypes observed in a patient (i.e., with a different phenotypes). After this step we obtained 36,041 negative (not causative) variant–disease pairs .

We downloaded phenotypes from the HPO database on 16 July 2020 and obtain 169,281 phenotype associations for 4,315 human genes. We downloaded phenotype associations from MGI on 20 March 2020 and obtained 228,214 associations between 13,529 mouse genes and classes in MP. We identify the human ortholog for mouse genes using the `HMD_HumanPhenotype.rpt` orthology file from MGI; the file contains 10,951 human orthologs for the 13,529 mouse genes resulting in 168,550 associations between human genes and MP classes. We obtain the GO annotations for human gene products from the GO Annotation database [2] on 20 March 2020 for 18,495 gene products with 495,719 annotations in total. We filtered out all the GO annotations with the evidence code indicating that the annotation was inferred from electronic annotation (IEA), or no biological data is available (ND). We map the UniProt accessions of the gene products to Entrez gene identifiers using the mappings provided by the Entrez database [3], and obtain 17,786 genes which have GO annotations for their gene product resulting in 208,630 associations between genes and GO classes. For the anatomical location of gene expression, we downloaded the GTEx Tissue Expression Profiles from the Gene Expression Atlas [4] which identifies gene expression across 53 tissues; 20,538 genes have an expression above the 4.0 threshold which was previously determined to be useful for predicting disease associations [5] resulting in 585,765 associations between genes and UBERON classes. We represent the tissues with their UBERON classes, excluding the tissues *transformed skin fibroblast* and *EBV-transformed lymphocyte* as these are not found in UBERON. For the cell type, we downloaded single-cell RNAseq data from the Tabula Muris project [6] in which genes are annotated with the CL. From this dataset, we obtain 6,559 human genes which have CL annotations, and 17,149 associations between genes and one or more classes from CL.

We use the combined PhenomeNET ontology [7] downloaded on 6 October, 2020, from the AberOWL ontology repository, as our phenotype ontology. PhenomeNET combines the phenotypes of human and other model organism as well as UBERON, GO and CL, and allows them to be compared.

### 2 Variant-based features

We use AnnotSV v2.3 [8], which uses data from multiple external databases to annotate and rank SVs based on the overlapping regions of the variants with known pathogenic variants in dbVar [9], the Database of Genomic Variants (DGV) [10], and disease-associated genes from OMIM. For each variant, AnnotSV generates annotations based on the variant length and the genes with which the variant overlaps (choosing among Refseq [11] gene annotations). Furthermore, AnnotSV reports the list of promoters with which the variant overlaps. From the annotations provided by AnnotSV, we use the variant length, variant type, GC content around the variant's breakpoints (GCcontent_left, GCcontent_right), and the number of promoters and genes affected by the variant as features.

We further use AnnotSV to obtain information about genes with which a structural variant overlaps: the length of the Coding DNA Sequence (CDS), transcript length (tx length), haploinsufficiency ranks collected from the Deciphering Developmental Disorders (DDD) study [12], haploinsufficiency (HI_DDDpercent) and triplosensitivity estimates from ClinGen [13], gene intolerance annotations from ExAC [14] including six annotations for synonymous variants (synZExAC), missense variants (misZExAC), loss of function variants (pLIExAC), deleation (delZExAC), duplications (dupZExAC), and CNV intolerance (cnvZExAC). Table 1 summarizes all the features used in our predictive model.

While not used as a feature of our prediction model, we also use AnnotSV to identify the allele frequency of variants using the 1,000 genomes allele frequency [15] and allele frequency from gnomAD [16]. We use this information to filter out common variants before applying our prediction.

### 3 Estimating variant pathogenicity by supervised prediction

#### 3.1 Phenotype prediction model

We apply DL2Vec to generate the feature representation for the patient phenotypes and genes. DL2Vec learns the "representation" for phenotypes and genes based on their annotations to ontology classes. The inputs to DL2Vec are associations of entities with ontology classes and the outputs are vectors (embeddings) of these entities. DL2Vec utilizes the axioms in ontologies to construct a graph representing phenotypes and their interrelations. DeepSVP incorporates biological background knowledge about the relation between phenotypes resulting from a loss of function in mouse/human genes, gene functions as defined using the GO, as well as the celltype and anatomical site of gene expression.

The phenotype model takes two vectors $v_1$ and $v_2$ as input, representing the embedding for the patient's phenotypes and the embedding for a gene, respectively. The embeddings are used as input for two neural network models $\mathsf{v}_1$ and $\mathsf{v}_2$. We then calculate the inner product for $\mathsf{v}_1(v_1)$ and $\mathsf{v}_2(v_2)$ and apply a sigmoid activation function to generate a prediction score, between the embedding for the phenotypes $v_1$, and gene $v_2$. We use binary cross-entropy as a loss function to train our model defined as:

$$Loss = -\frac{1}{N}\sum_{i=1}^{N} Y_i \cdot \log(P(Y_i)) + (1-Y_i)\cdot \log(1-P(Y_i)) \tag{1}$$

where $N$ correspond to the number of training samples, $Y_i$ is the true value for sample $i$, and $P(Y_i)$ is the predicted value for sample $i$.

Each neural network $\mathsf{v}_1$ and $\mathsf{v}_2$ consists of two hidden layers, in which the first layer with 256 units, and the second layer with 50 units. After each layer, we use dropout [17] with a rate of 20% , followed by a Leaky Rectified Linear Unit (LeakyReLU) [18] activation function. We use the Adam optimizer [19] to optimize the model parameters. We develop five different models using DL2Vec embeddings based on different feature types: functions of gene products (GO), mouse model phenotypes (MP), human phenotype (HP), celltype (CL), and site of expression (UBERON). For each set of phenotypes (characterizing a disorder, or the clinical phenotypes observed in an individual), and for each prediction model, we rank each gene based on the DL2Vec prediction score, from smallest to highest, and represent the association between them by their $m$-quantile in this distribution [20] (Figure 2 shows the distribution of the normalized quantile scores). This normalization and ranking aims to make prediction scores comparable across sets of phenotypes [21]. We use the quantile as one of the features of the combined prediction models.

#### 3.2 Combined prediction model

The combined prediction model uses the variant features and the phenotype-based scores produced by the DL2Vec-based predictions; the model is an artificial deep neural network model that uses genomic features derived from a variant as input together with the prediction score generated from the phenotype prediction model. We trained a separate model for each ontology dataset and aggregation type, either the maximum or average features scores for the genes within the variant region. The features used by the combined model are listed in the Supplementary Table 1.

Given a structural variant $\tau$ affecting regions that contain genes $G_1,...,G_n$, we obtain the phenotype prediction score $\phi(G_i)$ for the genes $G_1,...,G_n$ using each of the phenotype-based prediction models. We transform these scores into a feature for the variant $\tau$ using either the maximum or average of all the gene scores, i.e., either $\phi_{max}(\tau) = \max_{1\leq i\leq n}\phi(G_i)$ or $\phi_{avg}(\tau) = \frac{1}{n}\sum_{i=1}^{n}\phi(G_i)$. We normalize all the features using z-score normalization, in which the values for and feature $F$ are scaled based on the mean, and standard deviation of $F$. The value $v_i$ of $F$ is normalized to $\acute{v_i}$ by $\acute{v}_i = \frac{v_i-\mu_F}{\sigma_F}$ where $\acute{v_i}$ is z-score normalized one values, $v_i$ is the i-th value for the feature $A$, $\mu_F$ is the mean, and $\sigma_F$ the standard deviation, for feature $F$. We use the same mean and standard deviation to normalize the testing set.

We use 22 features for variants (8 features for the variant and 9 derived from the genes overlapping the variant) as well as 5 features from ontology embeddings. Some features are missing for some variants; to account for missing values, we use imputation. We imputed missing values by assigning them a zero value and additionally created indicator variables with a value set to 1 if the corresponding variant is missing and 0 otherwise. We use one-hot encoding to represent the categorical features with an "undefined" category for missing categorical annotations. We provide an analysis of feature importance and the correlation between features in the Supplementary Materials Section 3.3.

#### 3.3 Correlation between features

To test the redundancy of different scores of structural variants corresponding to the susceptibility of the phenotypes for the disease, we analyze the features by evaluating the pairwise correlation between them (Figure 1). According to their correlation coefficients, the features were broadly clustered into four major groups using the maximum and average score. As expected, measures of the structure of variants such as SV length and the number of genes or promoters were highly correlated. We further assess the importance of the features using two methods; the first using Extremely Randomized Trees ensemble learning Classifier (Extra Trees Classifier) (Figure 5), and the second using the Shapiro-

Wilk algorithm [22] (Figure 6). Both methods explore the linear relationships across features; Extra Trees Classifier aggregates the results of multiple decision trees and outputs the ranked features based on the information gain, while Shapiro-Wilk assesses the normality of the distribution of examples with respect to the feature. We noticed that the features rank using both methods are similar; however, both do not capture potential nonlinear relationships among features, so we included all the ranked features in our model.

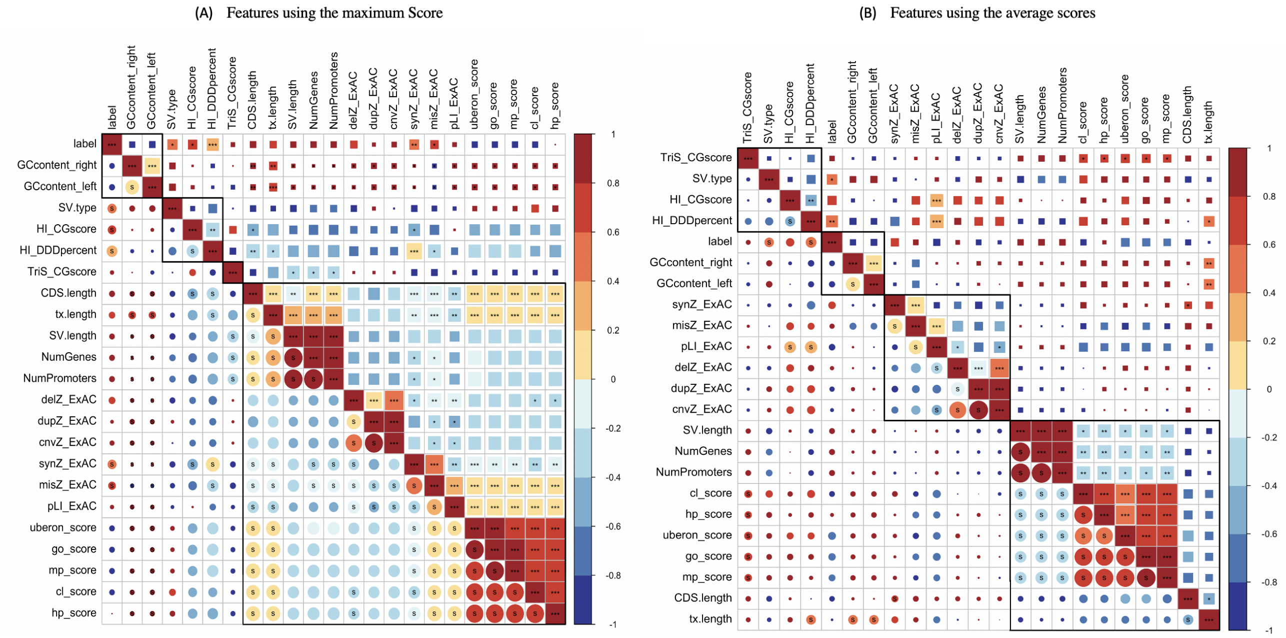


Figure 1: Correlation between the combined model features generated with *corrplot* package (version 0.84) in R, using *corrplot* function. 22 features of 42,202 SVs, (8 features for the variant, and 14 derived from the genes overlapping the variant) were obtained from the disease phenotypes and AnnotSV using various databases. The pairwise correlation was computed on all the features. Figure **A** shows results using the maximum score, and Figure **B** the average score. The features are ordered, and different clusters are highlighted based on the hierarchical clustering. Significant correlations ($P < 0.05$) are indicated by a letter 's' in the lower triangle. The color and size of circles represent the correlation strength (correlation coefficient). Statistical significance is indicated with asterisks ($^{*}$P <0.05;$^{**}$ P <0.01;$^{***}$ P <0.001).

### 4 Training and testing

We tuned for the following set of hyperparameters for the model: learning-rate $r \in [1 \times 10^{-6}, 1 \times 10^{-2}]$ with logarithmic transformation, and dropout $r \in [0.1, 0.5]$; number of layers $l \in [2, 6]$, and the number of nodes for each of the dense layers $n \in [50, 512]$; activation functions $a \in \{\text{relu}, \text{sigmoid}, \text{selu}\}$.

Following our experiments, the optimal parameters (learning rate, number of dense layers, number of dense nodes, dropout, activation) for each model were as follows: using the maximum score for the genes within the variant region, GO $(1 \times 10^{-3}, 6, 395, 0.126, \text{selu})$, MP $(1 \times 10^{-3}, 6, 214, 0.165, \text{sigmoid})$, HP $(1 \times 10^{-4}, 6, 391, 0.204, \text{selu})$, CL $(1 \times 10^{-5}, 2, 100, 0.2, \text{relu})$, UBERON $(2 \times 10^{-3}, 5, 414, 0.316, \text{selu})$, and combined model $(0.002, 6, 70, 0.132, \text{relu})$; and using the average score: GO $(9 \times 10^{-4}, 2, 147, 0.278, \text{relu})$, MP $(7 \times 10^{-5}, 3, 467, 0.121, \text{relu})$, HP $(1 \times 10^{-3}, 5, 121, 0.144, \text{relu})$, CL $(1 \times 10^{-3}, 3, 142, 0.235, \text{relu})$, UBERON $(3 \times 10^{-3}, 2, 420, 0.104, \text{sigmoid})$, and combined model $(3 \times 10^{-4}, 3, 439, 0.192, \text{relu})$. For all the models, we train for a maximum of 100 epochs. We use 15% of the training set for validation and stop training if the validation set loss increases for 10 epochs. We use a binary cross-entropy loss during training. We use a sigmoid activation function for the output layer.

### 5 Structural variant calling

We prepared 150-bp paired-end libraries using the TruSeq Nano DNA Sample Preparation kit (Illumina, USA). Sequencing was performed using an Illumina HiSeq 4000 at the Bioscience core laboratory, KAUST, with approximately 30X coverage. Following sequencing, we aligned reads to human genome build hg38 using the BWA MEM algorithm [23] and following the GATK standard workflows. We trim adapters using Trimmomatic (version 0.38), use BWA (version 0.7.17) for alignment, and samtools (version 1.8) [24] to remove duplicates and sort bam files. We use Manta (version 1.6) [25] to call structural variants. In total, we identified 8,723, 9,003, 9,608, 7,631, and 8,367 variants for the mother, father, first affected, second affected, and unaffected, respectively. We assume an autosomal recessive mode of inheritance or *de novo* variants, and use the pedigree to filter variants. We further filter common variants (minor allele frequency greater than 0.01) using gnomAD (version 2.1.1) [16] and the SVs from the 1000 genomes project. After filtering by pedigree, 148 structural variants remain; removing common variants reduced the number of variants to 47. We use the DeepSVP combined model with *maximum* as aggregation operation for the genes within the variant to prioritize disease-associated SVs.

| | Feature | Description |
|---|---|---|
| Structural Variant (SV) | SV length | Length of SV. |
| | SV type | Type of the SV (DEL, DUP) Categorical features. |
| | GCcontent_left* | Breakpoints annotations, GC content around the left SV breakpoint (+/- 100bp). |
| | GCcontent_right* | Breakpoints annotations, GC content around the right SV breakpoint (+/- 100bp). |
| | Number of genes* | Number of genes within the SVs. |
| | Number of promoters* | Number of promoters within the SVs. |
| | HI_CGscore | HaploInsufficiency Score (categorical features). |
| | TriS_CGscore | TriploSensitivity Score (categorical feature). |
| Genes-based annotations | DL2vec_Score* | Predict associations between genes and sets of phenotypes different ontolgies (GO, HP, CL, MP, and UBERON). |
| | CDS length* | Length of the CoDing Sequence (CDS) (bp) overlapping with the SV. |
| | tx length* | Length of transcript (bp) overlapping with the SV. |
| | synZ_ExAC* | Gene intolerance to synonymous variation. |
| | misZ_AxAC* | Gene intolerance to missense variation. |
| | pLI_ExAC* | The probability that a gene is intolerant to a loss of function variation. |
| | dupZ_ExAC* | Gene duplication intolerance. |
| | delZ_ExAC* | Gene deletion intolerance. |
| | cnvZ_ExAC* | Gne CNV intolerance. |
| | HI_DDDpercent* | Haploinsufficiency ranks, where in a single functional copy of a gene is insufficient to maintain normal function. |

Table 1: Annotation features for model training and prediction. An asterisk (*) indicates that a boolean indicator variable was created in order to handle undefined values for that feature.

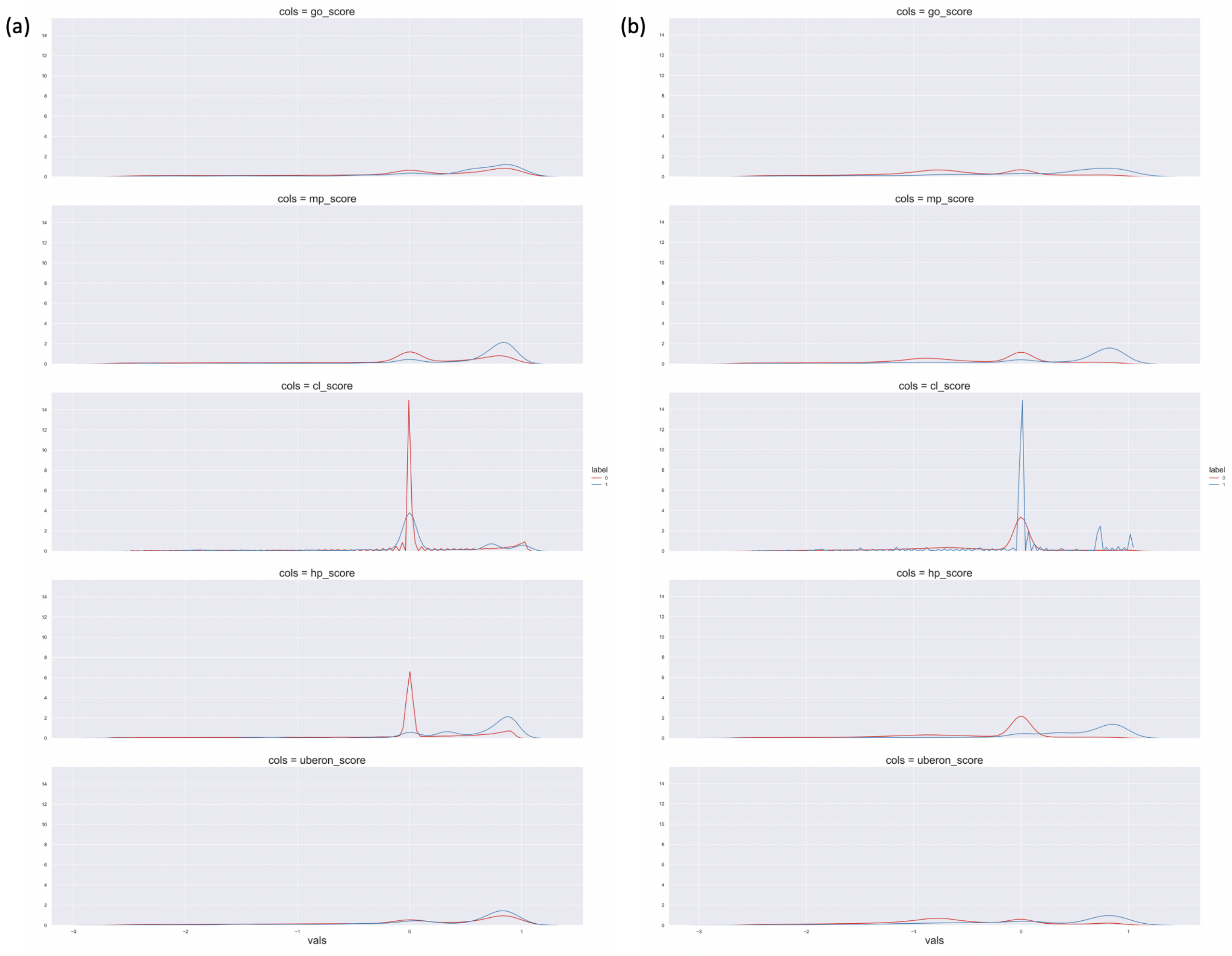


Figure 2: Distribution of the phenotype features, **(a)** using the maximum features scores, and **(b)** using the average score.

| **Aggregation method for genes within CNV** | **DeepSVP Models** | **ROCAUC** | **F1-Score** | **PRAUC** | **DOR** |
|---|---|---|---|---|---|
| **Maximum score** | GO | 0.9390 ± 0.0068 | 0.6681 ± 0.0127 | 0.8126 ± 0.0109 | 56.1942 ± 0.1290 |
| | MP | 0.9293 ± 0.0078 | 0.6900 ± 0.0134 | 0.7616 ± 0.0126 | 41.9207 ± 0.1075 |
| | HP | 0.9361 ± 0.0073 | 0.7009 ± 0.0129 | 0.7980 ± 0.0116 | 50.0758 ± 0.1132 |
| | CL | 0.9157 ± 0.0083 | 0.6394 ± 0.0134 | 0.7581 ± 0.0124 | 31.6137 ± 0.1067 |
| | UBERON | 0.9152 ± 0.0084 | 0.6011 ± 0.0137 | 0.7257 ± 0.0130 | 23.8252 ± 0.1027 |
| | GO+MP+HP+CL+UBERON | **0.9534 ± 0.0062** | **0.7397 ± 0.0125** | **0.8446 ± 0.0105** | **67.9658 ± 0.1177** |
| **Average score** | GO | 0.9186 ± 0.0081 | 0.6442 ± 0.0132 | 0.7189 ± 0.0127 | 35.4152 ± 0.1108 |
| | MP | 0.9336 ± 0.0074 | 0.6777 ± 0.0132 | 0.7801 ± 0.0120 | 41.3731 ± 0.1099 |
| | HP | 0.9373 ± 0.0072 | 0.7077 ± 0.0129 | 0.8287 ± 0.0110 | 51.1215 ± 0.1124 |
| | CL | 0.8913 ± 0.0093 | 0.5419 ± 0.0135 | 0.6656 ± 0.0134 | 17.3593 ± 0.1020 |
| | UBERON | 0.9040 ± 0.0089 | 0.5698 ± 0.0136 | 0.7029 ± 0.0132 | 20.0705 ± 0.1021 |
| | GO+MP+HP+CL+UBERON | **0.9505 ± 0.0062** | **0.7258 ± 0.0123** | **0.8520 ± 0.0101** | **73.3047 ± 0.1262** |

Table 2: Summary of the evaluation for predicting causative variants in our cross-validation testing data set

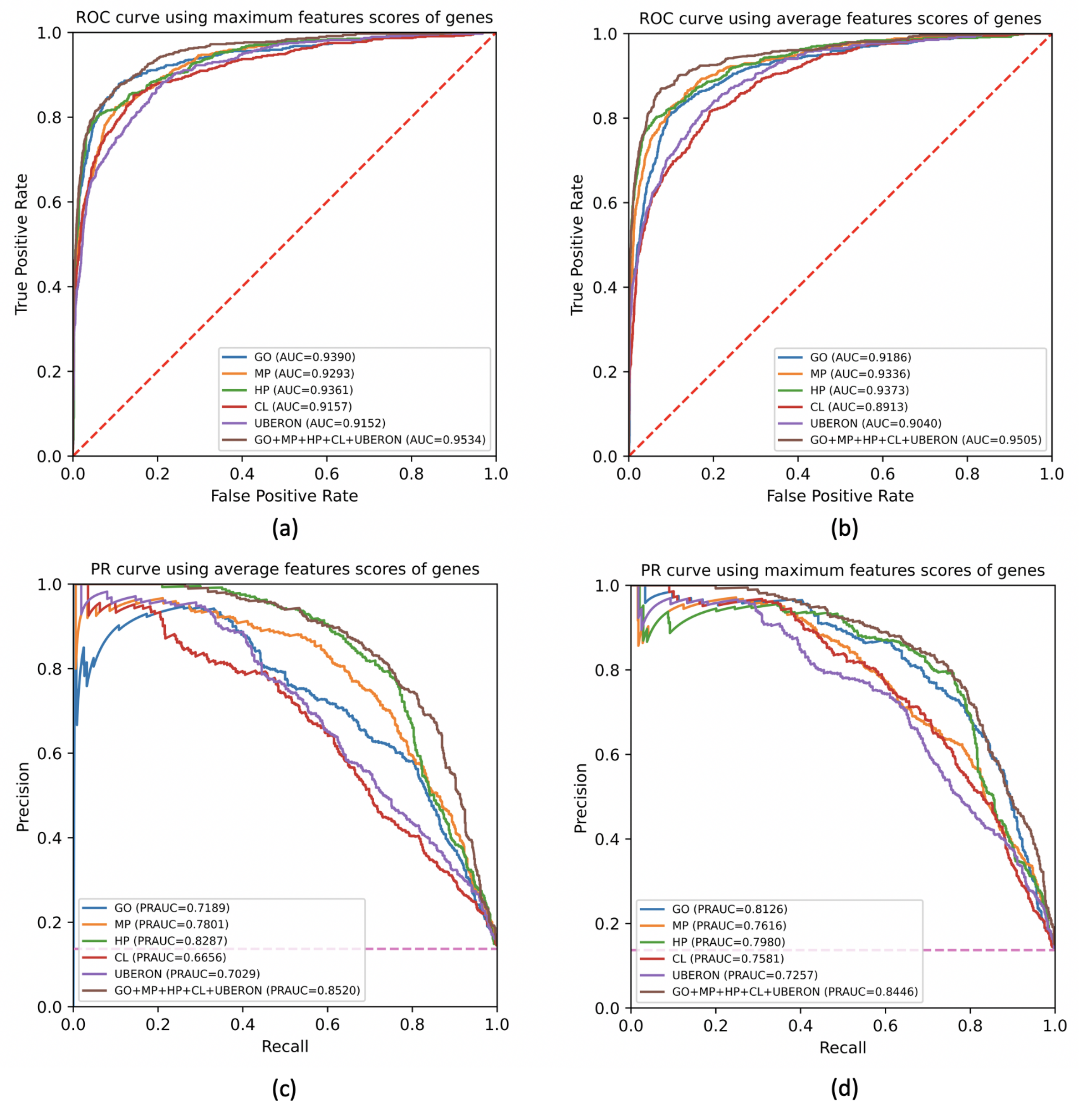


Figure 3: Summary of the ROCAUCs and PRAUCs curves for predicting causative variants using different features and ontologies, and different operation types. **(a)** the ROC curve using the maximum features scores; **(b)** the ROC curve using the average features scores; **(c)** the PR curve using the average features scores; and **(d)** the PR curve using the maximum features scores.

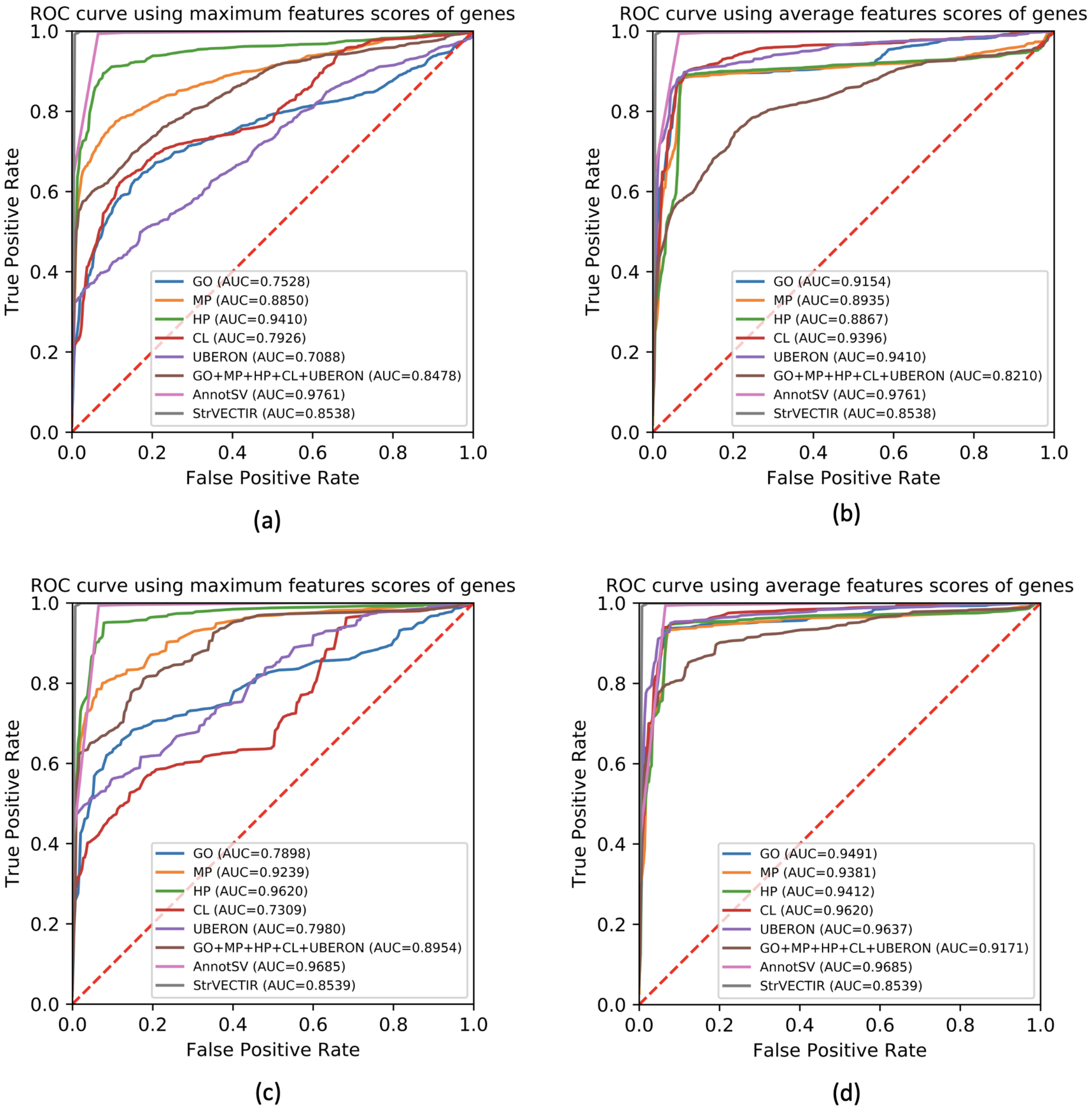


Figure 4: Summary of the ROCAUCs curves for the time-based split synthetic dataset using different ontologies, and different operation types in comparison to other methods. **(a)** and **(b)** for 1,503 newly added variants. **(c)** and **(d)** for 640 newly added variants associated with new diseases which were not present in our training dataset.

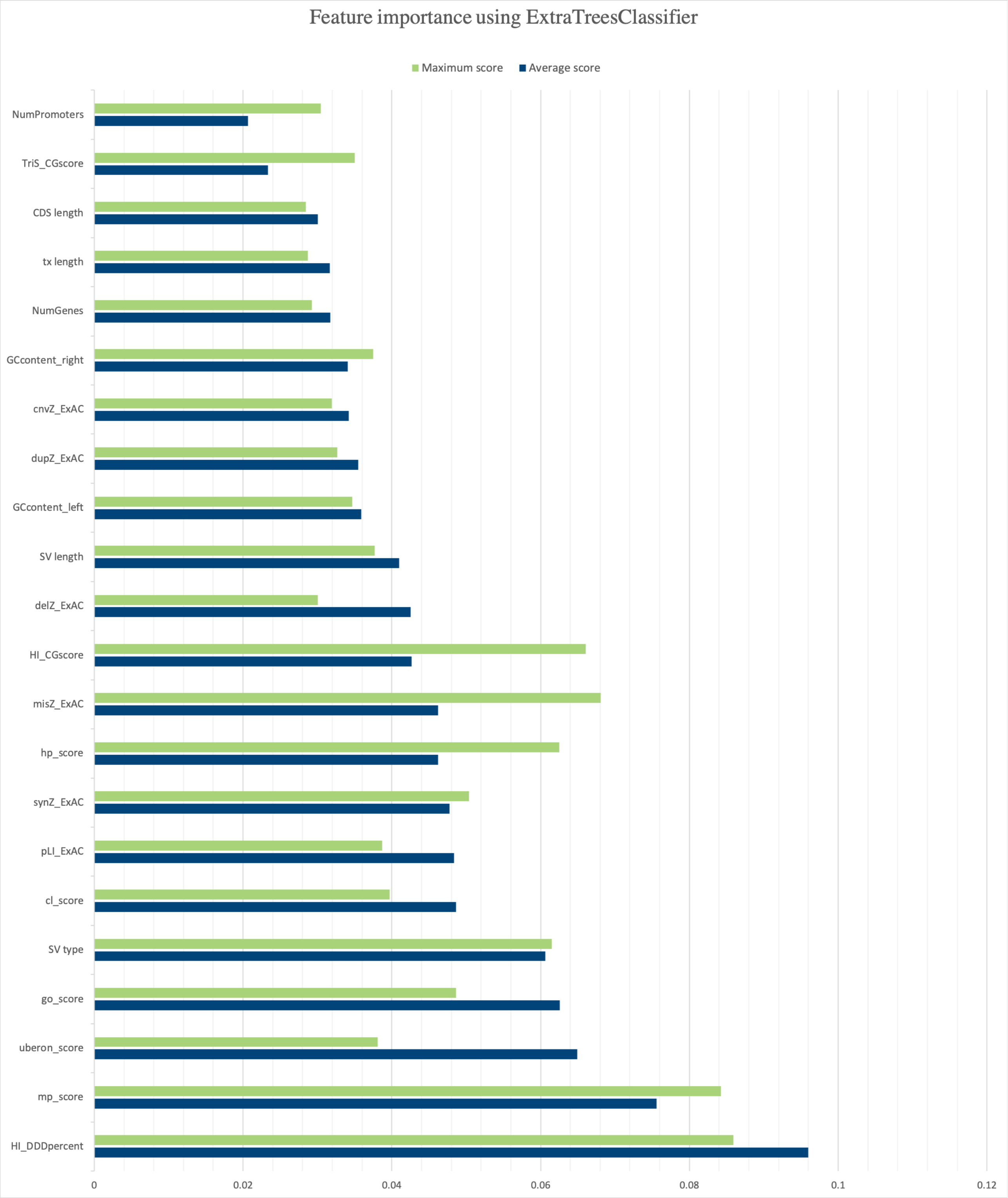


Figure 5: Feature importance using ensemble learning technique Extremely Randomized Trees Classifier (Extra Trees Classifier) that aggregates the results of multiple decision trees and output the ranked features based on the information gain.

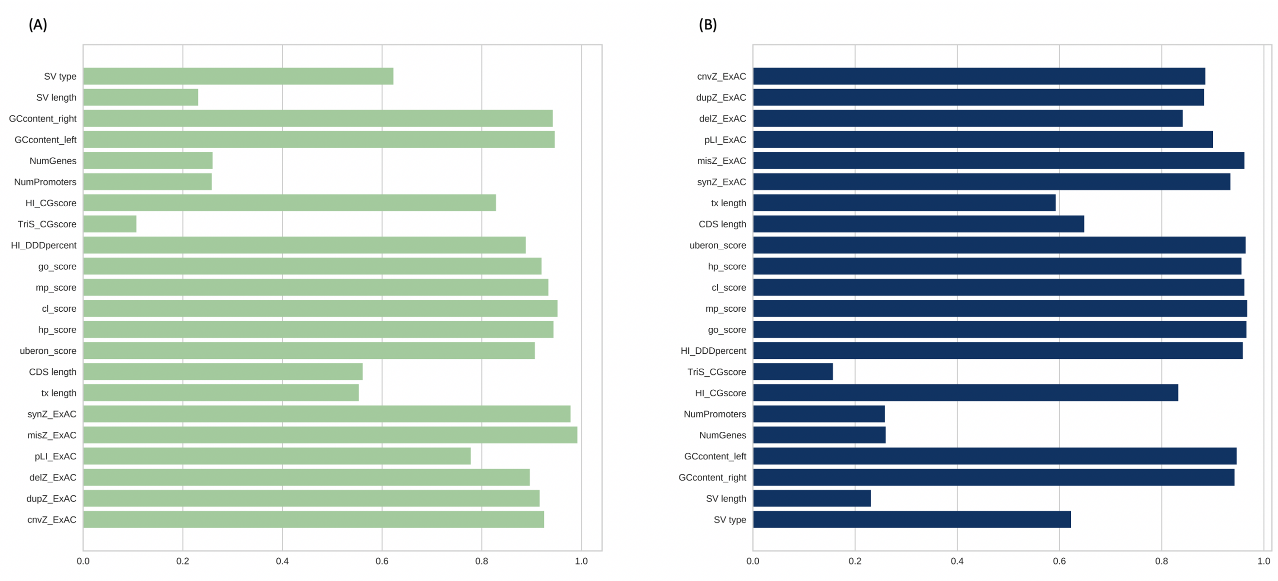


Figure 6: A one-dimensional ranking of features utilizing the Shapiro-Wilk algorithm generated using *Yellowbrick* Python package (version 1.0). The Shapiro algorithm takes into account a single feature at a time to assess the normality of the distribution of instances with respect to the feature. A barplot showing the relative ranks of each feature **(A)** using the maximum features scores, and **(B)** using the average score.

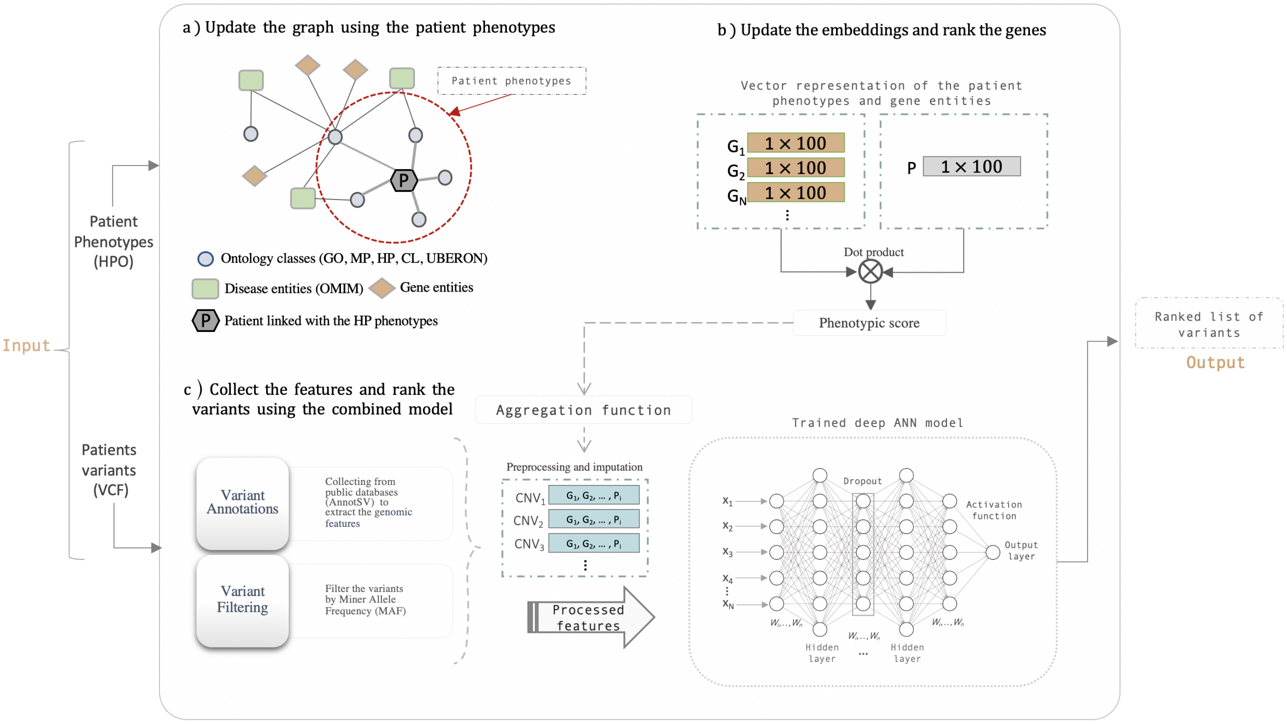


Figure 7: The workflow for DeepSVP inference. (**a**) Update the graph using the patient phenotypes, by adding a node for the patient and edges to the set of phenotypes. (**b**) Update the embeddings after generating the walks on the updated graph starting by the patient node, then run the DL2vec prediction model to rank the genes for the patient phenotypes using the patient and genes embeddings. (**c**) Collect the genomic features derived from the patient VCF and rank the variants using the DeepSVP combined prediction model, the rank determines how likely the variant is causative of the phenotypes observed in the patient. Abbreviations: `G`: Genes, `P`: Patient, `F`: Features, `P`: Phenotypic score.

| | | Synthetic dataset excluding 10% randomly of the phenotypes | | | | |
|---|---|---|---|---|---|---|
| **Aggregation method for genes within CNV** | **DeepSVP Models** | **Recall@1** | **Recall@10** | **Recall@30** | **ROCAUC** | **PRAUC** |
| Maximum score | GO | 27 (0.018) | 538 (0.358) | 826 (0.5496) | 0.7530 | 0.0242 |
| | MP | 65 (0.0432) | 985 (0.6554) | 1141 (0.7591) | 0.8854 | **0.0680** |
| | HP | 336 (0.2236) | 1093 (0.7272) | 1366 (0.9088) | 0.9403 | 0.0543 |
| | CL | 22 (0.0146) | 520 (0.346) | 873 (0.5808) | 0.7945 | 0.0299 |
| | UBERON | 74 (0.0492) | 528 (0.3513) | 624 (0.4152) | 0.7070 | 0.0339 |
| | GO+MP+HP+CL+UBERON | 173 (0.1151 | 834 (0.5549) | 917 (0.6101) | 0.8415 | 0.0471 |
| Average score | GO | 232 (0.1544) | 993 (0.6607) | 1332 (0.8862) | 0.9159 | 0.0629 |
| | MP | 27 (0.018) | 913 (0.6075) | 1333 (0.8869) | 0.8942 | 0.0642 |
| | HP | 74 (0.0492) | 668 (0.4444) | 1340 (0.8916) | 0.8880 | 0.0620 |
| | CL | 290 (0.1929) | 984 (0.6547) | 1360 (0.9049) | 0.9399 | 0.0379 |
| | UBERON | 488 (0.3247) | **1134 (0.7545)** | 1358 (0.9035) | 0.9414 | 0.0461 |
| | GO+MP+HP+CL+UBERON | 71 (0.0472) | 725 (0.4824) | 908 (0.6041) | 0.8202 | 0.0434 |
| SV pathogenicity prediction/classification | StrVCTVRE | 38 (0.0252) | 620 (0.4125) | 1022 (0.6799) | 0.8538 | 0.0270 |
| | AnnotSV | **930 (0.6188)** | 930 (0.6188) | **1493 (0.9933)** | **0.9760** | 0.0230 |

Table 3: Summary of the evaluation for predicting causative variants in the synthetic whole-genome benchmark dataset derived from dbVar, using 90% of the phenotypes. AnnotSV are computed based on ranking variants from "pathogenic" to"benign"; variants with the same classification are tied and we report all variants at the same (best) rank; Best performing results for each measure are indicated in bold.

| | | Synthetic dataset excluding 20% randomly of the phenotypes | | | | |
|---|---|---|---|---|---|---|
| **Aggregation method for genes within CNV** | **DeepSVP VModels** | **Recall@1** | **Recall@10** | **Recall@30** | **ROCAUC** | **PRAUC** |
| Maximum score | GO | 27 (0.018) | 538 (0.358) | 827 (0.5502) | 0.7533 | 0.0240 |
| | MP | 70 (0.0466) | 984 (0.6547) | 1146 (0.7625) | 0.8855 | **0.0674** |
| | HP | 332 (0.2209) | 1091 (0.7259) | 1371 (0.9122) | 0.9415 | 0.0550 |
| | CL | 23 (0.0153) | 519 (0.3453) | 870 (0.5788) | 0.7929 | 0.0298 |
| | UBERON | 74 (0.0492) | 527 (0.3506) | 625 (0.4158) | 0.7070 | 0.0340 |
| | GO+MP+HP+CL+UBERON | 167 (0.1111) | 866 (0.5762) | 949 (0.6314) | 0.8483 | 0.0505 |
| Average score | GO | 233 (0.155) | 993 (0.6607) | 1332 (0.8862) | 0.9158 | 0.0624 |
| | MP | 27 (0.018) | 912 (0.6068) | 1333 (0.8869) | 0.8942 | 0.0641 |
| | HP | 72 (0.0479) | 667 (0.4438) | 1342 (0.8929) | 0.8886 | 0.0630 |
| | CL | 290 (0.1929) | 988 (0.6574) | 1360 (0.9049) | 0.9397 | 0.0378 |
| | UBERON | 488 (0.3247) | **1133 (0.7538)** | 1358 (0.9035) | 0.9415 | 0.0460 |
| | GO+MP+HP+CL+UBERON | 71 (0.0472) | 725 (0.4824) | 908 (0.6041) | 0.8197 | 0.0436 |
| SV pathogenicity prediction/classification | StrVCTVRE | 38 (0.0252) | 620 (0.4125) | 1022 (0.6799) | 0.8538 | 0.0270 |
| | AnnotSV | **930 (0.6188)** | 930 (0.6188) | **1493 (0.9933)** | **0.9760** | 0.0230 |

Table 4: Summary of the evaluation for predicting causative variants in the synthetic whole-genome benchmark dataset derived from dbVar, using 80% of the phenotypes. AnnotSV are computed based on ranking variants from "pathogenic" to"benign"; variants with the same classification are tied and we report all variants at the same (best) rank; Best performing results for each measure are indicated in bold.

| | | Synthetic dataset excluding 30% randomly of the phenotypes | | | | |
|---|---|---|---|---|---|---|
| **Aggregation method for genes within CNV** | **DeepSVP Models** | **Recall@1** | **Recall@10** | **Recall@30** | **ROCAUC** | **PRAUC** |
| Maximumscore | GO | 27 (0.018) | 538 (0.358) | 827 (0.5502) | 0.7534 | 0.0242 |
| | MP | 67 (0.0446) | 983 (0.654) | 1139 (0.7578) | 0.8849 | **0.0673** |
| | HP | 334 (0.2222) | 1093 (0.7272) | 1373 (0.9135) | 0.9413 | 0.0548 |
| | CL | 23 (0.0153) | 519 (0.3453) | 871 (0.5795) | 0.7952 | 0.0299 |
| | UBERON | 75 (0.0499) | 529 (0.352) | 627 (0.4172) | 0.7079 | 0.0340 |
| | GO+MP+HP+CL+UBERON | 177 (0.1178) | 864 (0.5749) | 958 (0.6374) | 0.8483 | 0.0511 |
| Average score | GO | 234 (0.1557) | 990 (0.6587) | 1331 (0.8856) | 0.9157 | 0.0621 |
| | MP | 28 (0.0186) | 912 (0.6068) | 1333 (0.8869) | 0.8942 | 0.0646 |
| | HP | 72 (0.0479) | 677 (0.4504) | 1340 (0.8916) | 0.8887 | 0.0633 |
| | CL | 289 (0.1923) | 986 (0.656) | 1360 (0.9049) | 0.9397 | 0.0378 |
| | UBERON | 488 (0.3247) | **1133 (0.7538)** | 1358 (0.9035) | 0.9415 | 0.0458 |
| | GO+MP+HP+CL+UBERON | 73 (0.0486) | 728 (0.4844) | 910 (0.6055) | 0.8210 | 0.0432 |
| SV pathogenicity prediction/classification | StrVCTVRE | 38 (0.0252) | 620 (0.4125) | 1022 (0.6799) | 0.8538 | 0.0270 |
| | **AnnotSV** | **930 (0.6188)** | 930 (0.6188) | **1493 (0.9933)** | **0.9760** | 0.0230 |

Table 5: Summary of the evaluation for predicting causative variants in the synthetic whole-genome benchmark dataset derived from dbVar, using 70% of the phenotypes. AnnotSV are computed based on ranking variants from "pathogenic" to"benign"; variants with the same classification are tied and we report all variants at the same (best) rank; Best performing results for each measure are indicated in bold.

| | | Synthetic dataset excluding 50% randomly of the phenotypes | | | | |
|---|---|---|---|---|---|---|
| **Aggregation method for genes within CNV** | **DeepSVP Models** | **Recall@1** | **Recall@10** | **Recall@30** | **ROCAUC** | **PRAUC** |
| Maximum score | GO | 27 (0.018) | 538 (0.358) | 827 (0.5502) | 0.7543 | 0.0242 |
| | MP | 70 ( 0.0466 ) | 983 ( 0.654 ) | 1137 ( 0.7565 ) | 0.8847 | 0.0672 |
| | HP | 333 ( 0.2216 ) | 1092 ( 0.7265 ) | 1370 ( 0.9115 ) | 0.9407 | 0.0540 |
| | CL | 27 ( 0.018 ) | 538 ( 0.358 ) | 828 ( 0.5509 ) | 0.7931 | 0.0297 |
| | UBERON | 74 (0.0492) | 530 ( 0.3526 ) | 623 ( 0.4145 ) | 0.7071 | 0.0333 |
| | GO+MP+HP+CL+UBERON | 171 ( 0.1138 ) | 838 ( 0.5576 ) | 936 ( 0.6228 ) | 0.8423 | 0.0462 |
| Average score | GO | 235 ( 0.1564 ) | 992 ( 0.6600) | 1331 ( 0.8856 ) | 0.9157 | 0.0617 |
| | MP | 27 (0.018) | 915 ( 0.6088 ) | 1331 ( 0.8856 ) | 0.8940 | **0.0647** |
| | HP | 76 ( 0.0506 ) | 675 ( 0.4491 ) | 1333 ( 0.8869 ) | 0.8869 | 0.0611 |
| | CL | 288 ( 0.1916 ) | 987 ( 0.6567 ) | 1360 ( 0.9049 ) | 0.9398 | 0.0381 |
| | UBERON | 486 ( 0.3234 ) | **1133 (0.7538)** | 1358 (0.9035) | 0.9413 | 0.0458 |
| | GO+MP+HP+CL+UBERON | 70 ( 0.0466 ) | 725 ( 0.4824 ) | 910 ( 0.6055 ) | 0.8220 | 0.0412 |
| SV pathogenicity prediction/classification | StrVCTVRE | 38 (0.0252) | 620 (0.4125) | 1022 (0.6799) | 0.8538 | 0.0270 |
| | AnnotSV | **930 (0.6188)** | 930 (0.6188) | **1493 (0.9933)** | **0.9760** | 0.0230 |

Table 6: Summary of the evaluation for predicting causative variants in the synthetic whole-genome benchmark dataset derived from dbVar, using 50% of the phenotypes. AnnotSV are computed based on ranking variants from “pathogenic” to“benign”; variants with the same classification are tied and we report all variants at the same (best) rank; Best performing results for each measure are indicated in bold.

| | Rank using maximum scores | Rank using the average score |
|---|---|---|
| **GO** | 9 | 9 |
| **MP** | 7 | 10 |
| **HP** | 2 | 2 |
| **CL** | 9 | 12 |
| **UBERON** | 9 | 7 |
| **GO+MP+HP+CL+UBERON** | **1** | 5 |
| **StrVCTVRE** | 4 | |
| **AnnotSV** | **1** (pr:0.090) | |

Table 7: Ranking of disease-associated variant in a Saudi family using different DeepSVP models, and other methods. The ranks of the DeepSVP models are determined based on ranking a total of 47 variants. StrVCTVRE ranks only 6 out of 47 variants. AnnotSV predicted 11 variants as pathogenic out of 47 variants and we include the precision in parentheses.
